# Increasing equity in science requires better ethics training: a course by trainees, for trainees

**DOI:** 10.1101/2023.11.03.565577

**Authors:** Roshni A. Patel, Rachel A. Ungar, Alanna L. Pyke, Alvina Adimoelja, Meenakshi Chakraborty, Daniel J. Cotter, Malika Freund, Pagé Goddard, Justin Gomez-Stafford, Emily Greenwald, Emily Higgs, Naiomi Hunter, Tim M. G. MacKenzie, Anjali Narain, Daphne O. Martschenko

## Abstract

Despite the profound impacts of scientific research, few scientists have received the necessary training to productively discuss the ethical and societal implications of their work. To address this critical gap, we – a group of predominantly human genetics trainees – developed a course on genetics, ethics, and society. We intend for this course to serve as a template for other institutions and scientific disciplines. Our curriculum positions human genetics within its historical and societal context and encourages students to evaluate how societal norms and structures impact the conduct of scientific research. We demonstrate the utility of this course via surveys of enrolled students and provide resources and strategies for others hoping to teach a similar course. We conclude by arguing that if we are to work towards rectifying the inequities and injustices produced by our field, we must first learn to view our own research as impacting and being impacted by society.

## Introduction

Over the past few decades, rapid advancements in human genetics research have influenced many spheres of society, from reproductive health to criminal trials to tribal federal recognition claims.^1–3^ Moreover, discussions about genetic traits and ancestry testing are increasingly common in everyday conversations and popular culture.^4,5^ In the process, genetics has become deeply intertwined with political discourse – scientific arguments have been used to both combat and justify racist, classist, sexist, transphobic, and ableist ideologies. Though it has been long established that there is no genetic justification for racial classification,^6^ pseudoscientific analyses of genetics continue to proliferate across online forums, appearing to support white nationalist ideologies.^7,4,8^ Similarly, certain politicians have attempted to legislate gender and sexuality by leveraging arguments that incorrectly reference genetics, conflate sex and gender, and disregard the fact that sex determination is a biologically complex and nonbinary process.^9,10^

In response to the misuse of genetics research, ethicists and scientists alike have argued that scientists have a responsibility to society, not only in the questions they choose to ask, but also in how they communicate their findings.^11–16^ This sense of social responsibility particularly intensifies during a crisis when scientists are challenged to articulate their views on social and ethical issues.^17^ History is littered with examples of this ebb and flow.^11^ In the 1970s, several academics formed the Sociobiology Study Group and participated in the Committee Against Racism to counter the loud proponents of race- and sex-based genetic inferiority.^18,19^ More recently, the citing of genetics research by a white supremacist to justify his massacre of ten Black people in May 2022 served as a wakeup call for a number of researchers and funders to reflect on their impact on society.^16^

However, scientists have limited training in grappling with the social and ethical implications of their work. Trainees funded by the National Institutes of Health are required to receive education on the responsible conduct of research (RCR), but the majority of the RCR requirements are focused on research ethics internal to science. Research ethics programming predominantly evaluates scientists’ duties toward each other and spends little time interrogating the broader implications of scientific research within society.^20^ In addition, when institutional review boards (IRBs) evaluate human subjects research, they are – per the Common Rule – prohibited from considering any broad social or policy risks; they rarely, if ever, consider risks to anyone but research participants.^21–23^ As a result, the downstream implications of research, especially the social harms, are understudied and under-considered.^24^

Echoing other scholars, we argue that an essential part of our scientific training involves education on the intersection of science and society, as well as our responsibility to secure the ethical conduct and translation of our work.^14,16^ Across institutions, student-led advocacy has demonstrated the demand for more thoughtful post-graduate education on ethics and social responsibility.^25,26^ Integrating these topics into curricula has been shown to reap numerous benefits, including a decreased belief in racial essentialism, the perception that racial disparities and inequalities are primarily driven by genetics.^27,28^

In response to the gap we observed in our training, we – a trainee-led team affiliated with the Stanford Genetics Department – developed a course on genetics, ethics, and society. This course is intended to supplement the topics covered in the NIH RCR course. We successfully taught this course in Spring 2022 and 2023, and recently integrated it into the course recommendations for Stanford Genetics PhD students. Below, we describe our curriculum, evaluate its impacts on students, and suggest strategies for developing similar curricula at other universities. Our curriculum is available for others to use and adapt on our website.

### The course: Genetics, Ethics, and Society

Below, we briefly describe the background and development of our course, followed by an overview of the curriculum.

### Background and development

In the summer of 2020, during a Stanford Genetics town hall, PhD students, postdoctoral scholars, staff, and faculty co-developed a list of diversity, equity, and inclusion (DEI) initiatives to prioritize, one of which was increasing ethics education. Two of us (R.A.P. and R.A.U.) volunteered to lead this effort. After spending several months exploring different options for expanding ethics education, we decided that the best option would be to create a new course devoted to the intersection of genetics, ethics, and society.

We pitched our vision for the course to the Stanford McCoy Family Center for Ethics in Society, and received a grant to support curriculum development and teaching. After securing funding, we assembled a teaching team consisting of one bioethicist, one genetic counseling student, one genetic counselor, one genetics postdoc, and 7 genetics PhD students. The organizational structure consisted of one primary and secondary instructor for each class session, who were responsible for curriculum development; preparing lectures and activities; and teaching. In addition, two overall leads (R.A.P., R.A.U.) were responsible for administrative duties; managing the teaching team; designing course structure and learning goals; ensuring the cohesiveness across class sessions; providing feedback on curriculum development; and teaching. The leads additionally participated in the Course Design Institute hosted by the Stanford Center for Teaching and Learning and received feedback on curriculum design.^29^

The topics in our curriculum were chosen in part based on discussions with the broader genetics community, both from informal conversations as well as responses to an internal department survey of over 70 genetics PhD students. Our curriculum was also crafted around the specific expertise of our teaching team, which spanned a wide range of topics including forensic genetics, population genetics, bioethics, clinical genetic counseling, and genetic editing, as well as additional perspectives derived from lived experiences. We also leveraged external resources, such as Harvard Medical School’s Personal Genetics Education Project (pgEd, pged.org), a resource for the ethical, legal, and social implications of genetics. We iterated on curriculum development for six months before teaching our pilot course in 2022, which consisted of six two-hour classes taught over three weeks.

The following year, we expanded our course into ten two-hour classes that met once a week. Having demonstrated the success of our pilot course, we received funding directly from the Stanford Genetics Department to support our new teaching team: 5 genetics PhD students, 2 genetic counseling students, and one bioethicist. To ensure continuity, the two leads remained the same, two of the teaching staff were students from the 2022 course, and one was an instructor in the 2022 course. We worked on curriculum development for one quarter and revised the class based on student feedback from the previous year, conversations with faculty (D.O.M.), and discussions with the teaching team.

We had 20 students complete the course in 2022 and 22 students complete the course in 2023. Although the 2022 and 2023 courses are broadly similar, the second iteration allowed for more instructional time (20 hours vs 12 hours) and included new topics based on our 2022 student evaluations and the interests of the teaching team. In describing the curriculum below, we will focus on the 2023 extended course.

### Course overview

To develop our course from scratch, we utilized the backwards course design framework, in which the traditional order of designing a class is reversed by first identifying desired student outcomes, creating assessments that demonstrate student mastery of these outcomes, and then designing classroom lectures and activities to enable students to succeed in the assessments.^30^ Accordingly, we first envisioned our “big dream” for ethics education: that all biomedical scientists would have the tools to dissect the relationship between science and society; reflect on their own social responsibilities; and identify concrete actions towards a more equitable and socially responsible scientific enterprise.

We then identified three course-specific learning goals for students.^31^ First, students should be able to connect the historical context of genetics research to its modern-day practice. Given the American eugenics movement and its roots in California and Stanford in particular,^32,33^ we felt it was essential that Stanford students were familiar with this legacy of 20th century human genetics. Second, students should be able to analyze how societal norms and structures, along with personal identities, biases, and responsibilities, impact the conduct of scientific research. In other words, we wanted students to interrogate the perceived objectivity of science and assess how systems like racism, sexism, and colonialism impact(ed) the research questions that are proposed, funded, and published. Third, students should be able to evaluate the social and ethical implications of genetics research. We wanted students to both evaluate the impact of current research on society and anticipate the impact of future or proposed research by interrogating who benefits and who is harmed.

We scaffolded the content of each class session around class-specific learning objectives (Table 1). As the course progressed, these learning objectives increased in difficulty and complexity based on Bloom’s Taxonomy, a hierarchical pedagogical framework.^34^ We primarily achieved these learning objectives through a variety of active learning activities:^35^ each two-hour class typically featured short instructor-given lectures interspersed with student-led conversations where they analyzed case studies or role-played scenarios (Table 1).

**Table 1:** Overview of 2023 curriculum.

|  | Class Session | Content |
| --- | --- | --- |
| 1 | Course Introduction and History of Genetics | Learning objectives<br>• Define positionality, intersectionality, and privilege<br>• Understand how societal biases intersected with genetic research and medicine through the 20th century<br>Activities<br>• Students and instructors co-create ground norms for the classroom<br>• “Who’s in the room?” exercise: students anonymously describe identities they hold and reflect on the collection of identities in the classroom<br>• Privilege awareness exercise: instructors read aloud various social and economic privilege statements and students privately note which statements they identify with<br>• Reflection on positionality: students journal about which identities they hold, and how their identities impact the way students interact with the world |
| 2 | Introduction to Bioethics, Social Responsibility, and Community Engagement | Learning objectives<br>• Describe the principles of beneficence, non-maleficence, autonomy, and justice, and apply them to case studies<br>• Identify shortcomings of these four principles<br>• Understand the benefits of social responsibility and community-engaged research<br>• Identify key stakeholders and conflicts of stakeholdership in community-engaged research<br>Activities<br>• Analysis of *Havasupai Tribe v. the Arizona Board of Regents*: students identify stakeholders and discuss where and how conflicts arose between researchers and the Havasupai Tribe<br>• Role-play exercise: students are assigned roles as community members or researchers and co-design a research study using the principles of community-based participatory research |
| 3 | Race, Ancestry, and Genetics Part 1 | Learning objectives<br>• Distinguish between race, ethnicity, ancestry, and nationality<br>• Differentiate between social identity and genetic ancestry, and in particular, understand how social identity is (often) discretized, while genetic ancestry is a continuum<br>• Recall historical and societal issues that impact modern mistrust of medicine across certain racial and ethnic groups<br>• Understand how social determinants of health impact genetic medicine and research<br>Activities<br>• Word cloud exercise: students anonymously defined race, ethnicity, ancestry, and nationality, and reflected on the resulting word clouds<br>• Analysis of race and ancestry in healthcare: students discuss case examples and identify who benefitted, who was harmed, and how society was impacted more broadly |
| 4 | Race, Ancestry, and Genetics Part 2 | Learning objectives<br>• Describe societal perceptions of social identity (race, ethnicity, and nationality) and genetic ancestry, including racial essentialism<br>• Examine the consequences that these societal perceptions of social identity and genetic ancestry have on individuals and society broadly<br>• Critique scientific uses of social identity (race, ethnicity, and nationality) and genetic ancestry<br>Activities<br>• Analysis of popular media: students read a news article on genetic ancestry testing “surprises”, identify perceptions surrounding race and ancestry, and contrast genetic ancestry with social identities<br>• Analysis of scientific research: students read an academic paper, identify how the authors defined “populations” for their analysis, and discuss the impact on the interpretation of the paper<br>• Analysis of extremist views: students read a news article on white supremacist views of genetics and ancestry, identify misconceptions about race and ancestry, and discuss researchers’ responsibilities to mitigate harm in such contexts |
| 5 | Behavior and Genetic Determinism | Learning objectives<br>• Recognize how genetics research can be mis-used to justify genetic determinist viewpoints and the associated harms<br>• Describe current efforts by the research community to combat the weaponization of research, and articulate strategies that individual scientists can employ<br>Activities<br>• Analysis of genetic determinism: students read articles on the genetic basis of educational attainment, athleticism, or sexuality, and discuss societal implications of this research<br>• Analysis of FAQ utility: students read a sociobehavioral genetics paper, brainstorm questions, and evaluate whether their questions were successfully answered in the FAQ accompanying the paper<br>• Reflection on social responsibility: students discuss whether researchers have a responsibility to combat the harms of genetic determinism and brainstorm potential actions |
| 6 | Sex and Gender; Ability | Learning objectives<br>• Distinguish the relationship between biological sex, gender identity, and sexuality through discussion of the role genetics plays in these concepts<br>• Examine the medicalization of identities and how it impacts society, medicine, and research<br>• Critique genetics studies and practices from a disability-rights perspective<br>Activities<br>• Word cloud exercise: students anonymously defined sex, gender, and sexuality, and reflected on the resulting word clouds<br>• Role-play exercise: students role-play as a governing sports body conducting “sex verification” tests for athletes and discuss the challenges in making these decisions and their impacts on athletes |
| 7 | Screening and Selection | Learning objectives |
|  |  | • Understand the role of eugenics in the development of the prenatal genetic testing field<br>• Summarize the current landscape of prenatal genetic testing, carrier screening, embryo selection and gene editing<br>• Extrapolate the current and future implications of the currently available technologies with a focus on reproductive choice and disability rights<br>• Evaluate how ethics can help guide society towards ethical application of these technologies |
|  |  | Activities<br>• Reflection on eugenics: students draw connections between the eugenics movement and reproductive health<br>• Analysis of current technologies: students discuss individual motivations and broader ethical implications at stake in contemporary case studies of prenatal screening<br>• Analysis of potential future technologies: students evaluate pre-implantation genetic testing and germline genetic engineering through the lens of each of the four principles of bioethics |
| 8 | Identification and Privacy | Learning objectives<br>• Analyze the role that government, private, and public databases play in the application of forensic genetics in the criminal justice system globally<br>• Examine risks and benefits of identification and privacy from private and public entities |
|  |  | Activities<br>• Analysis of forensic genetics: students read case studies on the usage of forensic genetics and assess the benefits and harms of using genetic information in each instance<br>• Analysis of data policies: students read privacy and consent policies for various consumer and research databases, and evaluate how well the benefits and harms are communicated to participants<br>• Reflection on social responsibility: students brainstorm potential harms posed by participants in genetic databases, and the social responsibilities of data stewards |
| 9 | Genetics in the Archives | Learning objectives<br>• Understand how individuals acted to counter the misuse of genetics research to further racist and sexist ideologies<br>• Connect historical examples of ethical problems to the modern-day case studies presented in this course<br>• Analyze how scientists' ideologies impacted their interpretation of science |
|  |  | Activities<br>• Analysis of primary source documents: students read and discuss primary source documents curated around 5 different topics; see Supplementary Material for details on topics and documents |
| 10 | Self-Reflection and Action | Learning objectives<br>• Evaluate your own positionality and its relationship to your work as a scientist<br>• Construct and reflect on viewpoints that recognise and relate to |
|  |  | different positionalities<br>• Ideate ways to make existing academic structures more equitable, inclusive, and just; create actionable goals to do so |
|  |  | Activities<br>• Reflection on the course: students note down their main takeaways from each class session on post-it notes around the classroom<br>• Reflection on reflexivity: students revisit their reflection on positionality and journal about how their positionality impacts their work as a researcher<br>• Reflection on post-course action: students independently brainstorm their “big dream” for science and identify actions they can take towards this goal over different timescales and different personal/professional spheres |

Importantly, we aimed to not make our course prescriptive. We instead focused on giving students the tools to reason through ethical problems themselves. To do so, we tried to create an environment in which people with varied perspectives could express their thoughts. When possible, we also exposed students to multiple viewpoints on a single issue and reiterated the complexity and nuance in ethical case studies.

Below, we summarize our curriculum. We additionally detail the learning objectives and active learning activities for each class in Table 1, and provide the full curriculum on our website.

### Description of curriculum

In the first class, we established an open and inclusive environment by co-creating classroom norms and introducing the concepts of identity, positionality, intersectionality, and privilege. Next, we delved into the history of genetics, because we believe that deconstructing the myth of scientific objectivity is easiest to understand through historical examples. We traced the impact of societal influences on genetics from the Classical Era to modern times, emphasizing its relevance through local and global examples of eugenic practices and their lasting legacies.^33,36–39^ Following this discussion of history, we covered the origins of bioethics, introduced key ethical principles and guidelines such as the Belmont Report, and prompted students to consider the social responsibilities of researchers towards both research subjects and society more broadly.^40,14,41^ In doing so, students explored the limitations of existing ethical guidelines and societal protections, as well as the role that community engagement and community-based participatory research play in tackling this problem.^22,42,43^

After students developed a foundation for the history of human genetics and bioethics, we turned to understanding the relationship between race, ancestry, and genetics.^44,45^ Students examined how misconceptions can manifest in both scientific research and clinical practice, and discussed the downstream implications for various aspects of society, including perceptions of self-identity and racial health disparities.^46–48^ Next, students explored the role that contemporary genetics research plays in beliefs about genetic and racial essentialism, and considered the responsibility of scientists in averting such social harms of research.^49,50^ We continued our examination of the relationship between genetics and identity by exploring the medicalization of identities, specifically in the context of gender, sexuality, and (dis)ability.^9,51,52^ Students then discussed ethical implications in the current landscape of reproductive and forensic genetics, as well as potential future developments in the field.^1,53,54,3^

For the penultimate class, we brought students to the Stanford library to directly engage with historical documents. To choose the documents, we (R.A.P. and R.A.U.) initially surveyed hundreds of materials in the Stanford University Archives that pertained to genetics, ethics, and society, and ultimately curated a selection of documents relating to five topics: historical perspectives on eugenics; historical perspectives on race, genetics, and IQ; sociobiology and scientific activism; community activism against studying biological causes of violence; and community activism against forced sterilization (details in Supplemental Material). For each topic, students spent some time looking through the documents before we facilitated group discussions about the content. After spotlighting these historical examples of scientific and community-led activism, we dedicated the last class to self-reflection: we asked students to interrogate their own roles and responsibilities as scientists and imagine how to achieve their vision of a more equitable and just scientific enterprise.

## Results

To assess the impact of our course on students, we offered an optional and anonymous pre- and post-course survey in both 2022 and 2023. We had 19 total respondents in 2022 (15 pre-course survey, 14 post-course survey), and 30 total respondents in 2023 (26 pre-course survey, 15 post-course survey). Pre- and post-course survey data could not be linked for enrolled students who missed one of the two surveys; students who dropped the course after taking the first survey; or students who forgot their survey identification code.

### Student demographics

Course participants ranged from undergraduates to postdoctoral scholars, but the course was mainly composed of early-stage graduate students (54%; Supplemental Figure 1A). The majority of students identified as women (61%; Supplemental Figure 1B). Moreover, the majority of students also reported belonging to additional identities that are underrepresented in STEM (61%; Supplemental Figure 1C) (including but not limited to identities around race/ethnicity, socioeconomic status, sexuality, ability, and family education levels). These course demographics paralleled the demographics of course instructors. This is consistent with previous literature demonstrating underrepresented students spend more time on advocacy and diversity efforts and are more likely to teach about ethical or societal issues.^55–57^

### Student evaluations

Students’ motivations for taking this course varied greatly, but were primarily to learn about current ethical issues and to develop the tools to discuss them (Supplemental Figure 2). For multiple students, a central motivation for taking the course was to gain a better understanding of the use of ancestry and race in genetics research; they were concerned about “the risk of categorizing race and ethnic backgrounds in studies that can be misconstrued”. Geneticists can be uninformed and imprecise in their use of race and ancestry; numerous studies have observed that genetics researchers use population labels in highly variable and unsystematic ways.^58–60^ At best, this creates ambiguity for the reader; at worst, it muddles these concepts and promotes racial essentialism.

Through survey data and conversations with students, we observed that in 2022, students left the course still struggling to understand the concepts of genetic ancestry and race (Supplemental Figure 4A). As a result, we doubled the amount of time spent on this concept in 2023 and reworked the section by borrowing heavily from the recently published National Academies report, “Using Population Descriptors in Genetics and Genomics Research”.^48^ In doing so, we discussed both the risks and benefits associated with the use of race and ancestry in genetic research, giving students the tools to independently critique their usage and evaluate the societal implications. By the end of the course in 2023, all students agreed there are risks associated with considering race or genetic ancestry in research, and students were divided on whether there are benefits (Supplemental Figure 4D, 4F). One student said they were “empowered to continue to learn about the use of race/ethnicity information in genetic studies and to improve my influence in this area as a human genetics researcher”.

Consistent with previous work,^61^ we found that perceptions of scientific objectivity can be influenced by classroom interventions. At the end of both classes, we found that students are better able to reason through the relationship between science and society: all students agreed with the statement: “scientific research is influenced by societal norms, structures, and values” (Supplemental Figure 3). One student remarked that “genetics and society can’t be separated from each other”. In addition, students were able to incorporate historical context into their understanding of the relationship between science and society: students reported strong gains in understanding the influence of the eugenics movement on modern-day genetics research (Figure 1A). One student stated “the foundation of the field I work in was based in eugenics, and I need to recognize how those thoughts and ideas have pervaded current scientific understanding”, exemplifying an awareness of scientific reflexivity.

**Figure 1:**
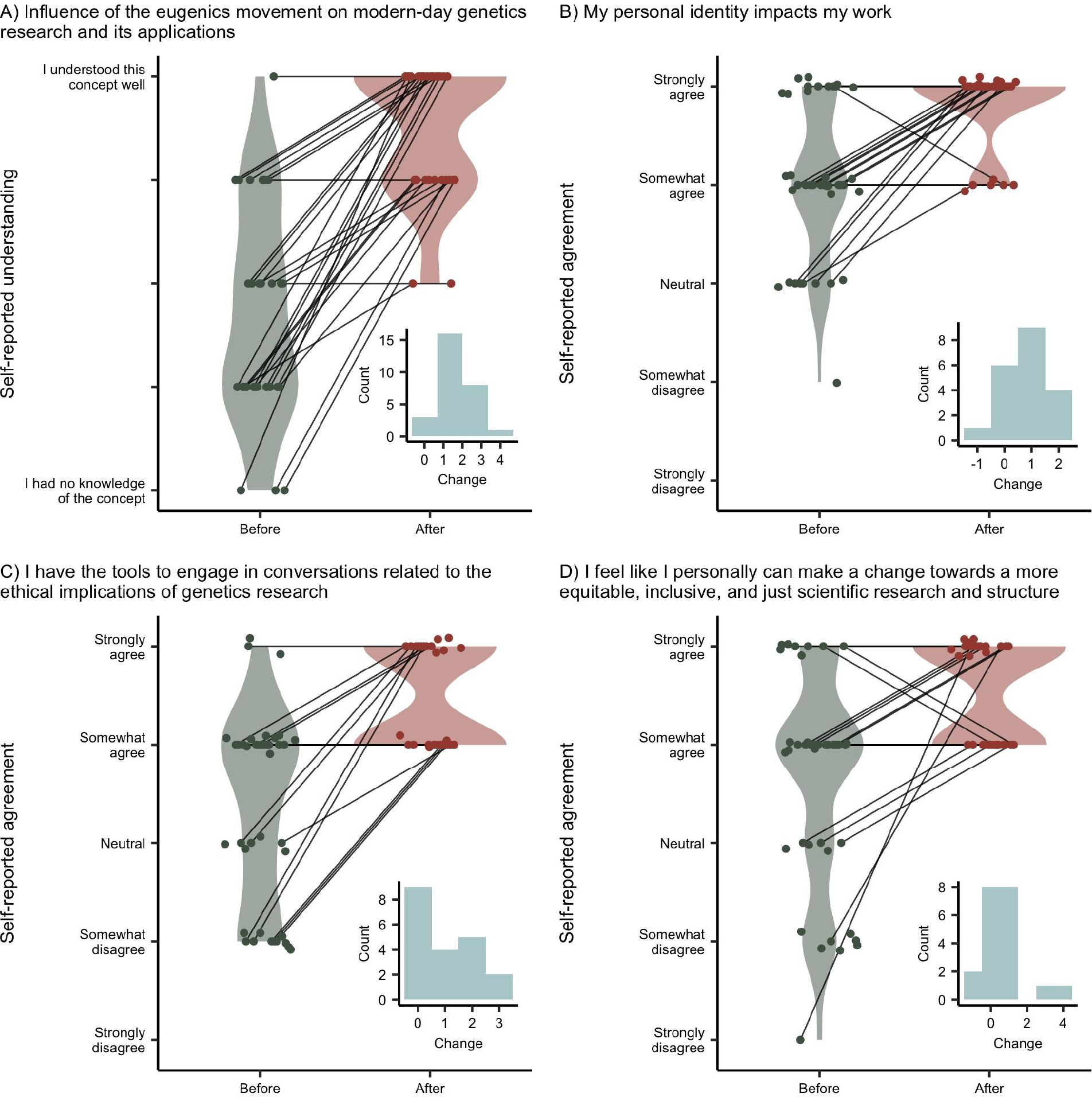
Key insights from student evaluations. A) Change in self-reported understanding of the “influence of the eugenics movement on modern-day genetics research and its applications” before and after the course. B) Change in self-reported agreement with the statement “my personal identity impacts my work” before and after the course. C) Change in self-reported agreement with “I have the tools to engage in conversations related to the ethical implications of genetics research” before and after the course. D) Change in self-reported agreement with “I feel like I personally can make a change towards a more equitable, inclusive, and just scientific research and structure” before and after the course. Data points are aggregated between 2022 and 2023 and jittered for enhanced visibility. Violin plots are included for illustrative purposes, with solid lines connecting the pre-course and post-course responses from the same student when applicable. Plot insets illustrate the change in opinion between pre-course and post-course responses for students who took both surveys, where a value of one represents a 1-unit difference in understanding or agreement.

Most importantly, students were able to apply this same critical lens to their own work. Previous work has shown that over 80% of life scientists agreed that research has ethical and societal impacts, but fewer than 30% believed their own research had such impacts.^62,63^ Consequently, a central objective of our course was to enable students to reflect on the social impacts of their research and their responsibilities to mitigate against potentially harmful impacts. By the end of the course in both 2022 and 2023, all students agreed that their personal identity impacts their work (Figure 1B). One student stated that their main takeaway was the “importance of thinking about the ethical implications of one’s work”, and another noted their takeaway was “consider[ing] the benefits and harm of my work, who are the stakeholders and who benefits and participates in my work”. Moreover, about half of students expressed increased agreement with the statements “I have the tools to engage in conversations related to the ethical implications of genetics research” and “I feel like I personally can make a change towards a more equitable, inclusive, and just scientific research and structure” (Figure 1C, D). One student said they will take away “how to be a better advocate for more ethical science”, and another wrote “science is deeply flawed and rooted in racism. It is our responsibility as scientists to speak out against it”, demonstrating a strengthened awareness and agency in relation to their own social responsibilities.

### Lessons learned

A class on genetics, ethics, and society naturally results in difficult conversations, and requires vulnerability, empathy, and trust from its students. We spent considerable time on building classroom norms (e.g. being respectful of conflicting viewpoints and committing to learning and growing) so that students felt comfortable voicing disagreements with each other and with the instructors, both of which are essential to student learning. Throughout the class, we received positive feedback about the classroom environment. Several students also commented that they preferred small group discussions over discussions with the entire class, and that power structures within small groups (e.g. graduate students vs postdocs) impacted their comfort and ease of sharing. For others looking to facilitate similar discussions, we suggest taking the time to create a supportive classroom environment, and to do so by prioritizing small group discussions between peers.

Consistent with a wealth of academic literature,^64–67^ active learning activities were particularly highly regarded by students. A standout example was a role-play activity in which students were randomly assigned different stakeholder roles in co-designing a research study using principles from community-engaged research. In general, we noticed during role-play activities that students were highly attentive and felt strongly about the roles they adopted; this is consistent with studies suggesting role-play increases ethical sensitivity.^68^

In order to make our classes on the history of genetics feel more impactful, we deliberately focused on Stanford and the Bay Area, and had students directly interact with primary source documents, which has been shown to foster a deeper understanding of historical contexts.^69,70^ For example, students were able to page through a 1973 article in the Stanford Daily describing a debate between several Stanford professors on the alleged genetic basis of racial differences in intelligence.^71^ Due to our focus on local events and hands-on learning, students shared that they felt “closer” to the history. For others teaching similar classes, we believe emphasizing these local connections and examples would be beneficial, as would interacting with primary source documents when possible.

We consistently observed that students benefited from the synergy between class topics. For example, students frequently cited the class on genetics and disability during their discussions of reproductive genetics the following week. Likewise, our discussions of race and ancestry were an important precursor to the classes on genetic determinism and privacy and identification. We encourage other course developers to carefully consider the myriad connections between seemingly disparate topics, and to leverage them to maximize students’ synthesis.

We believe students felt empowered, rather than defeated, at the end of the course because throughout the course, we intentionally reiterated that students were active participants in the scientific structure. This was especially emphasized during our visit to the Stanford Archives, which highlighted historical examples of community- and scientist-led activism. Students were surprised and impressed, and mentioned they felt there was less scientific activism happening today. As a result, we found that these powerful examples of historical activism were a well-placed segue into our final class encouraging our students to envision what changes they might start to make in their own scientific communities. We intentionally saved the most pointed self-reflection for the end of the course given the amount of knowledge, humility, and unlearning necessary to interrogate one’s own role and responsibility as a scientist.

In surveys and personal conversations, students provided several suggestions for areas of improvement. For each class, we usually had at least 40 cumulative minutes devoted to discussion time, but in both 2022 and 2023, students reported that there was not enough time for discussion. Moreover, students also said that they would appreciate greater depth in material. For any course, there is a trade-off between breadth and depth, and we often struggled with this balance. To ensure an effective and tailored learning experience, course developers should consider their own resources as well as the needs of their potential students when deciding whether to create a shorter survey course or a longer course with more in-depth content.

## Discussion

In short, we developed a course on genetics, ethics, and society to tackle a significant gap in graduate student education. Elsewhere, ethics courses have been seen as unhelpful “hoops to jump through” or “too philosophical”, even by scientists who care about ethics.^72,73^ In contrast, we found that not only was our course generally enjoyed by students, it was also successful in accomplishing its learning goals. This is consistent with previous work which has shown that emphasizing practicality (e.g. by focusing on tangible societal impacts) results in more effective ethics courses.^74^

We note an important limitation of this course: the material is targeted for a primarily American audience that does human genetics research. However, a course of this nature can be modified for other audiences and objectives, and we encourage others to use this course framework and adapt it in response to the current moment, the expertise of the teaching team, the focus of their department, the local geographic and historical context, and the interests of the students in the course. Finally, additional topics could be covered: other topics requested by students included science policy, science communication, the role of industry, non-human genetics, and funding structures.

It should be acknowledged that this was a profound learning experience for us as a teaching team. Advocacy efforts for DEI often help trainees develop valuable skills that are not otherwise learned in the laboratory – we not only gained skills in pedagogy but also in effective collaboration, leadership, management, and communication. Burnout is a major issue in DEI efforts, and studies have shown burnout is linked to feeling undervalued and poorly compensated.^75^ We strongly believe that this course only materialized because our labor was compensated, first through an internal grant and later through department funding. Moreover, the funding structure of our institution and the support of our advisors allowed us to dedicate the time to develop and teach this course.

As with most trainee-led initiatives, long-term sustainability is always a challenge. Early on, we (R.A.P. and R.A.U.) sought to promote the longevity of our course by securing long-term funding from our department such that in every future year, TAs will be compensated for curriculum development and teaching. We were also intentional about the composition of the teaching team: in both 2022 and 2023, the teaching team had several early-stage PhD students, guaranteeing continuity between years when Masters students or late-stage PhD students graduate. Moreover, we had frequent and open discussions with TAs about leadership transitions to ensure a smooth transition when our tenure concluded.

Ultimately, we have illustrated an effective ethics course that enables students to critically reflect on the interplay of science and society, and their own role in the scientific structure. In their own words, students said they took away that “the field of genetics has a history of eugenics and this has shaped what has been studied up until this point”, and that “current sociopolitical climate and norms influence the type of research/questions that people are interested in answering”.

They emphasized it was important to “ensur[e] that genetics and all biomedical research does not cause further harm to already marginalized communities”. By the end of the class, they felt like they gained the tools to “articulate these conversations with other scientists and also those that do not know science” and “be a better advocate for more ethical science”, as well as the “importance of thinking about the ethical implications of one[‘]s work”.

We hope to enable others to build similar courses at other institutions and to facilitate this, we have made our entire curriculum available on our [website](https://stanford-genethics.github.io/). We believe that education on social responsibility and the ethical implications of scientific research is a necessary step towards building a more equitable and just scientific enterprise.

## Ethical considerations

Before administering the survey, we received approval from the Stanford Internal Review Board (IRB protocol 655032). The surveys were administered by an individual who was not affiliated with the class. The survey administrator followed a pre-approved script to inform students of the potential harms and benefits of filling out the survey, after which they were given 15 minutes of class time to fill out the survey. All instructors left the room during this time. Additionally, the survey stayed active for 24 hours to ensure that even the survey administrator would not know if a student chose to not fill out the survey. Students were asked to use a randomly generated ID they were randomly handed to enable us to link pre- and post-course surveys.Only the survey administrator had access to these codes, and the protocol director anonymized these codes prior to allowing others access to the data. To further ensure there were no consequences for students, grades for the class were only determined by attendance, and survey results were not accessed by course instructors until after the grading period was over.

## Acknowledgements

We are extraordinarily thankful for Julie Baker, who provided crucial guidance, was an incredible advocate, and generally uplifted us during this process. This course would not have been possible without funding from the Stanford Genetics Department, The Genetics Advocacy Committee at Stanford (TGAC), and most importantly, the Ethics, Society, and Technology Hub in the Stanford McCoy Family Center for Ethics in Society. We are indebted to Josh Tycko who led the grant application process, along with many other TGAC members who contributed towards it. We want to thank Ashlyn Jaeger and Wendy Christiansen for their key administrative support throughout the course. We want to thank Hank Greely, Hua Tang, Kelly Ormond, Stan Cohen, Abbey Thompson, and results from an anonymous survey of Stanford Neuroscience PhD students for advice when deciding how to proceed with increasing ethics education in the department, as well as Mildred Cho who gave advice throughout. We appreciate review of the manuscript that led to improvements from Tami Gjorgjieva, Mildred Cho, Stephen Montgomery, Abbey Thompson, Kelly Ormond, and Callie Chappell. This course benefited immensely from knowledge gained in the Graduate Course Design Institute, and we utilized the Stanford Center for Teaching and Learning during this design process. Devon Bonner provided invaluable guidance in the process of writing and submitting an IRB. We are appreciative of Jazlyn Mooney, Alice Popejoy, Ronit Mazzoni, and Marc Feldman for providing guest lectures in our class, all of which we later heard from students were really helpful in their learning and understanding. We want to thank the staff with Stanford Special Collections, especially Regina Roberts and Tim Noakes, who enabled our class in the Stanford Archives and contributed so much to our own learning. Thank you to all of our PIs for providing us the space and grace to take the time to develop and teach this course. We also acknowledge the Muwekma Ohlone community, the traditional stewards and caretakers of the land that Stanford is on, and the land on which we developed and taught this course. Finally, we are immensely grateful to our students who trusted us in leading and designing this course, and for the students who took the time to give feedback to ensure we can improve the class going forward and share this information beyond Stanford.

## Notes

### Competing Interest Statement

The authors have declared no competing interest.

## References

1. St. John, P. (2020). The untold story of how the Golden State Killer was found: A covert operation and private DNA. Los Angel. Times.

2. Severson, A.L., Byrd, B.F., Mallott, E.K., Owings, A.C., DeGiorgio, M., de Flamingh, A., Nijmeh, C., Arellano, M.V., Leventhal, A., Rosenberg, N.A., et al. (2022). Ancient and modern genomics of the Ohlone Indigenous population of California. Proc. Natl. Acad. Sci. 119, e2111533119. 10.1073/pnas.2111533119.

3. Kirksey, E. (2023). Opinion | Does Gene Editing Have a Future in Reproductive Medicine? N. Y. Times.

4. Mittos, A., Zannettou, S., Blackburn, J., and Cristofaro, E.D. (2020). “And We Will Fight for Our Race!” A Measurement Study of Genetic Testing Conversations on Reddit and 4chan. Proc. Int. AAAI Conf. Web Soc. Media 14, 452–463. 10.1609/icwsm.v14i1.7314.

5. Ogbunugafor, C.B., and Edge, M.D. (2022). Gattaca as a lens on contemporary genetics: marking 25 years into the film’s “not-too-distant” future. Genetics 222, iyac142. 10.1093/genetics/iyac142.

6. Lewontin, R.C. (1972). The Apportionment of Human Diversity. In Evolutionary Biology: Volume 6, T. Dobzhansky, M. K. Hecht, and W. C. Steere, eds. (Springer US), pp. 381–398. 10.1007/978-1-4684-9063-3_14.

7. Price, M. (2018). ‘It’s a toxic place.’ How the online world of white nationalists distorts population genetics. ScienceInsider.

8. Carlson, J., and Harris, K. (2022). The apportionment of citations: a scientometric analysis of Lewontin 1972. Philos. Trans. R. Soc. B Biol. Sci. 377, 20200409. 10.1098/rstb.2020.0409.

9. 9. Ainsworth, C. (2015). Sex redefined. Nature 518, 288–291. 10.1038/518288a.

10. Hassan, A. (2023). States Passed a Record Number of Transgender Laws. Here’s What They Say. N. Y. Times.

11. Beckwith, J. (1993). A Historical View of Social Responsibility in Genetics. BioScience 43, 327–333. 10.2307/1312065.

12. Stern, J.E., and Elliot, D. (1997). The Ethics of Scientific Research (University Press of New England).

13. Gibbons, M. (1999). Science’s new social contract with society. Nature 402, C81–84. 10.1038/35011576.

14. Beckwith, J., and Huang, F. (2005). Should we make a fuss? A case for social responsibility in science. Nat. Biotechnol. 23, 1479–1480. 10.1038/nbt1205-1479.

15. Martschenko, D.O., and Smith, M. (2021). Genes do not operate in a vacuum, and neither should our research. Nat. Genet. 53, 255–256. 10.1038/s41588-021-00802-5.

16. Carlson, J., Henn, B.M., Al-Hindi, D.R., and Ramachandran, S. (2022). Counter the weaponization of genetics research by extremists. Nature 610, 444–447. 10.1038/d41586-022-03252-z.

17. Ludmerer, K.M. (1969). American geneticists and the eugenics movement: 1905–1935. J. Hist. Biol. 2, 337–362. 10.1007/BF00125023.

18. Jon Beckwith and Bob Lange (1978). AAAS: Sociobiology on the Run. Sci. People 10, 38–39.

19. Freda Salzman (1979). Sociobiology: The Controversy Continues. Sci. People 11, 20–27.

20. National Institutes of Health, Agency for Healthcare Research and Quality, and Health Resources & Services Administration (2022). Updated Guidance: Requirement for Instruction in the Responsible Conduct of Research.

21. Emanuel, E.J., Wendler, D., and Grady, C. (2000). What Makes Clinical Research Ethical? JAMA 283, 2701–2711. 10.1001/jama.283.20.2701.

22. Klitzman, R.L. (2013). How IRBs View and Make Decisions About Social Risks. J. Empir. Res. Hum. Res. Ethics JERHRE 8, 58–65. 10.1525/jer.2013.8.3.58.

23. The Federal Policy for the Protection of Human Subjects (2018).

24. Trejo, S., Wedow, R., and Martschenko, D.O. (2022). Scientists Must Consider the Risk of Racist Misappropriation of Research. Sci. Am. https://www.scientificamerican.com/article/scientists-must-consider-the-risk-of-racist-misappropriation-of-research/.

25. Hall, T.E., Engebretson, J., O’Rourke, M., Piso, Z., Whyte, K., and Valles, S. (2017). The Need for Social Ethics in Interdisciplinary Environmental Science Graduate Programs: Results from a Nation-Wide Survey in the United States. Sci. Eng. Ethics 23, 565–588. 10.1007/s11948-016-9775-0.

26. Green, K.-A., Wolinsky, R., Parnell, S.J., del Campo, D., Nathan, A.S., Garg, P.S., Kaplan, S.E., and Dasgupta, S. (2022). Deconstructing Racism, Hierarchy, and Power in Medical Education: Guiding Principles on Inclusive Curriculum Design. Acad. Med. 97, 804. 10.1097/ACM.0000000000004531.

27. Donovan, B.M. (2022). Ending genetic essentialism through genetics education. Hum. Genet. Genomics Adv. 3, 100058. 10.1016/j.xhgg.2021.100058.

28. Lynn, T.M., D’urzo, K.A., Vaughan-Ogunlusi, O., Wiesendanger, K., Colbert-Kaip, S., Capcara, A., Chen, S., Sreenan, S., and Brennan, M.P. (2023). The impact of a student-led anti-racism programme on medical students’ perceptions and awareness of racial bias in medicine and confidence to advocate against racism. Med. Educ. Online 28, 2176802. 10.1080/10872981.2023.2176802.

29. Modell, A. (2021). Teacher-centered vs. Student-centered course design. In.

30. Bowen, R.S. (2017). Understanding by Design. Vanderbilt Univ. Cent. Teach. https://cft.vanderbilt.edu/understanding-by-design/.

31. Simon, B., and Taylor, J. (2009). What is the Value of Course-Specific Learning Goals?

32. Stanford Eugenics History Project Eugen. Stanf. https://www.stanfordeugenics.com.

33. Stern, A.M. (2015). Eugenic Nation: Faults and Frontiers of Better Breeding in Modern America 2nd ed.

34. Krathwohl, D.R. (2002). A Revision of Bloom’s Taxonomy: An Overview. Theory Pract. 41, 212–218. 10.1207/s15430421tip4104_2.

35. Baepler, P., Walker, J., Brooks, D., Saichaie, K., and Petersen, C. (2016). A Guide to Teaching in the Active Learning Classroom: History, Research, and Practice 10.4324/9781003442820.

36. Zimmer, C. (2018). She Has Her Mother’s Laugh: The Powers, Perversions, and Potential of Heredity (Penguin).

37. Charmantier, I. (2020). Linnaeus and Race. Linn. Soc. https://www.linnean.org/learning/who-was-linnaeus/linnaeus-and-race.

38. Jindia, S. (2020). Belly of the Beast: California’s dark history of forced sterilizations. The Guardian.

39. Arce, J. (2021). The long history of forced sterilization of Latinas. UnidosUS. https://unidosus.org/blog/2021/12/16/the-long-history-of-forced-sterilization-of-latinas/.

40. The National Commission for the Protection of Human Subjects of Biomedical and Behavioral Research (1979). Belmont Report: Ethical Principles and Guidelines for the Protection of Human Subjects of Research (United States Department of Health, Education and Welfare).

41. Resnik, D.B., and Elliott, K.C. (2016). The Ethical Challenges of Socially Responsible Science. Account. Res. 23, 31–46. 10.1080/08989621.2014.1002608.

42. Claw, K.G., Anderson, M.Z., Begay, R.L., Tsosie, K.S., Fox, K., and Garrison, N.A. (2018). A framework for enhancing ethical genomic research with Indigenous communities. Nat. Commun. 9, 2957. 10.1038/s41467-018-05188-3.

43. Lemke, A.A., Esplin, E.D., Goldenberg, A.J., Gonzaga-Jauregui, C., Hanchard, N.A., Harris-Wai, J., Ideozu, J.E., Isasi, R., Landstrom, A.P., Prince, A.E.R., et al. (2022). Addressing underrepresentation in genomics research through community engagement. Am. J. Hum. Genet. 109, 1563–1571. 10.1016/j.ajhg.2022.08.005.

44. Coop, G. (2019). Reading tea leaves? Polygenic scores and differences in traits among groups. Preprint at arXiv, 10.48550/arXiv.1909.0089210.48550/arXiv.1909.00892.

45. Mathieson, I., and Scally, A. (2020). What is ancestry? PLOS Genet. 16, e1008624. 10.1371/journal.pgen.1008624.

46. Harmon, A. (2018). Why White Supremacists Are Chugging Milk (and Why Geneticists Are Alarmed). N. Y. Times.

47. Padawer, R. (2018). Sigrid Johnson Was Black. A DNA Test Said She Wasn’t. N. Y. Times Mag.

48. Using Population Descriptors in Genetics and Genomics Research: A New Framework for an Evolving Field (2023). (National Academies Press) 10.17226/26902.

49. Martschenko, D.O., Domingue, B.W., Matthews, L.J., and Trejo, S. (2021). FoGS provides a public FAQ repository for social and behavioral genomic discoveries. Nat. Genet. 53, 1272–1274. 10.1038/s41588-021-00929-5.

50. Riskin, J., and Feldman, M.W. (2022). Why Biology Is Not Destiny. N. Y. Rev. Books 69.

51. Maxmen, A. (2019). Controversial ‘gay gene’ app provokes fears of a genetic Wild West. Nature 574, 609–610. 10.1038/d41586-019-03282-0.

52. Mukherjee, D., Tarsney, P.S., and Kirschner, K.L. (2022). If Not Now, Then When? Taking Disability Seriously in Bioethics. Hastings Cent. Rep. 52, 37–48. 10.1002/hast.1385.

53. Barnert, E., Katsanis, S.H., Mishori, R., Wagner, J.K., Selden, R.F., Madden, D., Berger, D., Erlich, H., Hampton, K., Kleiser, A., et al. (2021). Using DNA to reunify separated migrant families. Science 372, 1154–1156. 10.1126/science.abh3979.

54. Turley, P., Meyer, M.N., Wang, N., Cesarini, D., Hammonds, E., Martin, A.R., Neale, B.M., Rehm, H.L., Wilkins-Haug, L., Benjamin, D.J., et al. (2021). Problems with Using Polygenic Scores to Select Embryos. N. Engl. J. Med. 385, 78–86. 10.1056/NEJMsr2105065.

55. Bielefeldt, A.R., Polmear, M., Knight, D., Swan, C., and Canney, N. (2018). Intersections between Engineering Ethics and Diversity Issues in Engineering Education. J. Prof. Issues Eng. Educ. Pract. 144, 04017017. 10.1061/(ASCE)EI.1943-5541.0000360.

56. Bielefeldt, A.R., Lewis, J.W., Polmear, M., Knight, D., and Swan, C. (2021). Alumni Reflect on Their Education About Ethical and Societal Issues. In.

57. Kamceva, M., Kyerematen, B., Spigner, S., Bunting, S., Li-Sauerwine, S., Yee, J., and Gisondi, M. (2022). More Work, Less Reward: The Minority Tax on US Medical Students. J. Wellness 4. 10.55504/2578-9333.1116.

58. Hunt, L.M., and Megyesi, M.S. (2008). Genes, Race and Research Ethics: Who’s Minding the Store? J. Med. Ethics 34, 495–500. 10.1136/jme.2007.021295.

59. Panofsky, A., and Bliss, C. (2017). Ambiguity and Scientific Authority: Population Classification in Genomic Science. Am. Sociol. Rev. 82, 59–87. 10.1177/0003122416685812.

60. Duello, T.M., Rivedal, S., Wickland, C., and Weller, A. (2021). Race and genetics versus ‘race’ in genetics. Evol. Med. Public Health 9, 232–245. 10.1093/emph/eoab018.

61. Williams, C.T., and Rudge, D.W. (2016). Emphasizing the History of Genetics in an Explicit and Reflective Approach to Teaching the Nature of Science. Sci. Educ. 25, 407–427. 10.1007/s11191-016-9821-y.

62. McCormick, J.B., Boyce, A.M., and Cho, M.K. (2009). Biomedical Scientists’ Perceptions of Ethical and Social Implications: Is There a Role for Research Ethics Consultation? PLoS ONE 4, e4659. 10.1371/journal.pone.0004659.

63. McCormick, J.B., Boyce, A.M., Ladd, J.M., and Cho, M. (2012). Barriers to Considering Ethical and Societal Implications of Research: Perceptions of Life Scientists. AJOB Prim. Res. 3, 40–50. 10.1080/21507716.2012.680651.

64. Deslauriers, L., Schelew, E., and Wieman, C. (2011). Improved Learning in a Large-Enrollment Physics Class. Science 332, 862–864. 10.1126/science.1201783.

65. Freeman, S., Eddy, S.L., McDonough, M., Smith, M.K., Okoroafor, N., Jordt, H., and Wenderoth, M.P. (2014). Active learning increases student performance in science, engineering, and mathematics. Proc. Natl. Acad. Sci. 111, 8410–8415. 10.1073/pnas.1319030111.

66. Tharayil, S., Borrego, M., Prince, M., Nguyen, K.A., Shekhar, P., Finelli, C.J., and Waters, C. (2018). Strategies to mitigate student resistance to active learning. Int. J. STEM Educ. 5, 7. 10.1186/s40594-018-0102-y.

67. Theobald, E.J., Hill, M.J., Tran, E., Agrawal, S., Arroyo, E.N., Behling, S., Chambwe, N., Cintrón, D.L., Cooper, J.D., Dunster, G., et al. (2020). Active learning narrows achievement gaps for underrepresented students in undergraduate science, technology, engineering, and math. Proc. Natl. Acad. Sci. 117, 6476–6483. 10.1073/pnas.1916903117.

68. Jasemi, M., Goli, R., Zabihi, R.E., and Khalkhali, H. (2022). Educating ethics codes by lecture or role-play; which one improves nursing students’ ethical sensitivity and ethical performance more? A quasi-experimental study. J. Prof. Nurs. Off. J. Am. Assoc. Coll. Nurs. 40, 122–129. 10.1016/j.profnurs.2021.11.002.

69. American Historical Association Committee of Seven (1909). The Study of History in Schools: Report to the American Historical Association by the Committee of Seven (Macmillan).

70. Barton, K.C., and Levstik, L.S. (2004). Teaching History for the Common Good (Routledge).

71. Kershner, V., and Jacobs, J. (1973). Cavalli-Sforza, Shockley, X Debate Tonight. Stanf. Dly.

72. Smith-Doerr, L. (2008). Decoupling Policy and Practice: How Life Scientists Respond to Ethics Education. Minerva 46, 1–16. 10.1007/s11024-007-9084-5.

73. Brosnan, C., and Cribb, A. (2014). Between the bench, the bedside and the office: The need to build bridges between working neuroscientists and ethicists. Clin. Ethics 9, 113–119. 10.1177/1477750914558549.

74. Katsarov, J., Andorno, R., Krom, A., and van den Hoven, M. (2022). Effective Strategies for Research Integrity Training—a Meta-analysis. Educ. Psychol. Rev. 34, 935–955. 10.1007/s10648-021-09630-9.

75. Verduzco-Gutierrez, M., Larson, A.R., Capizzi, A.N., Bean, A.C., Zafonte, R.D., Odonkor, C.A., Bosques, G., and Silver, J.K. (2021). How Physician Compensation and Education Debt Affects Financial Stress and Burnout: A Survey Study of Women in Physical Medicine and Rehabilitation. PM&R 13, 836–844. 10.1002/pmrj.12534.

