## Supplemental for "Increasing equity in science requires better ethics training: a course by trainees, for trainees"

### Supplementary Material

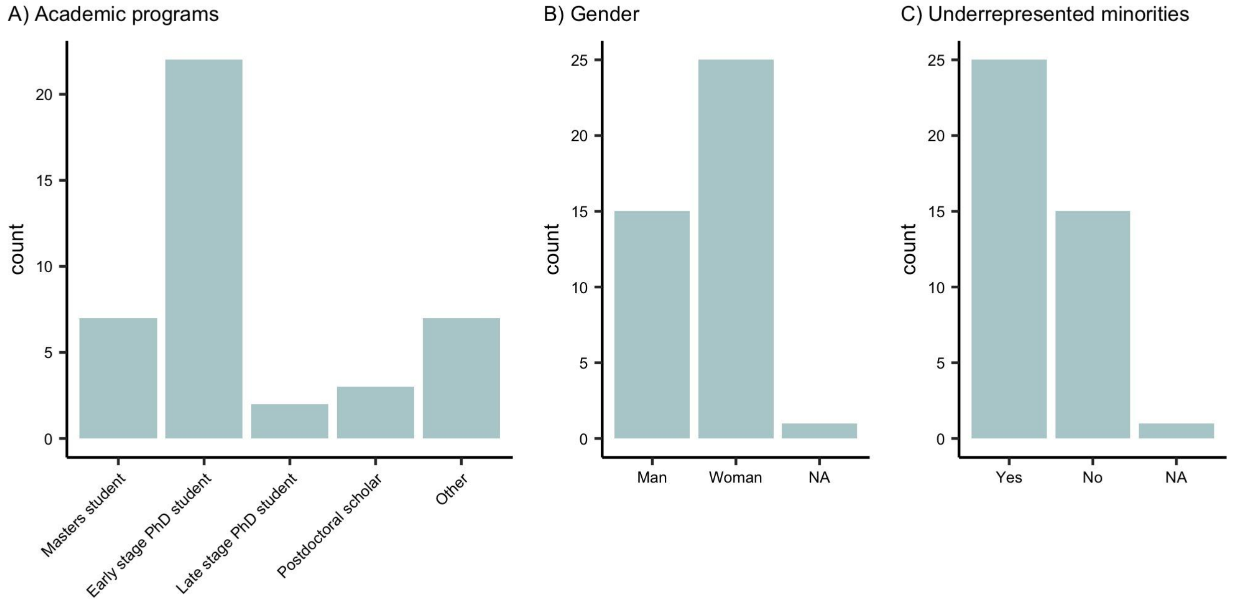


**Supplemental Figure 1: Demographics of enrolled students.** Histograms depict self-reported A) academic programs, B) gender, and C) identification as an underrepresented minority in STEM. Data points are aggregated between 2022 and 2023. For A), "Early stage PhD student" refers to years 1-3, and "Late stage PhD student" refers to years 4+. "Other" consists of students who were undergraduates or staff, or did not respond (NA). For B), students were asked if they identify as a man, woman, non-binary, and/or transgender, with the ability to select multiple answers. For C), students were asked "Aside from gender, do you otherwise identify as being part of a historically excluded or otherwise marginalized population? (e.g. Black, Indigenous, Hispanic/Latino, Hawaiian/Pacific Islanders, low-income students, first-generation college students, LGBTQ+, students with disabilities)".

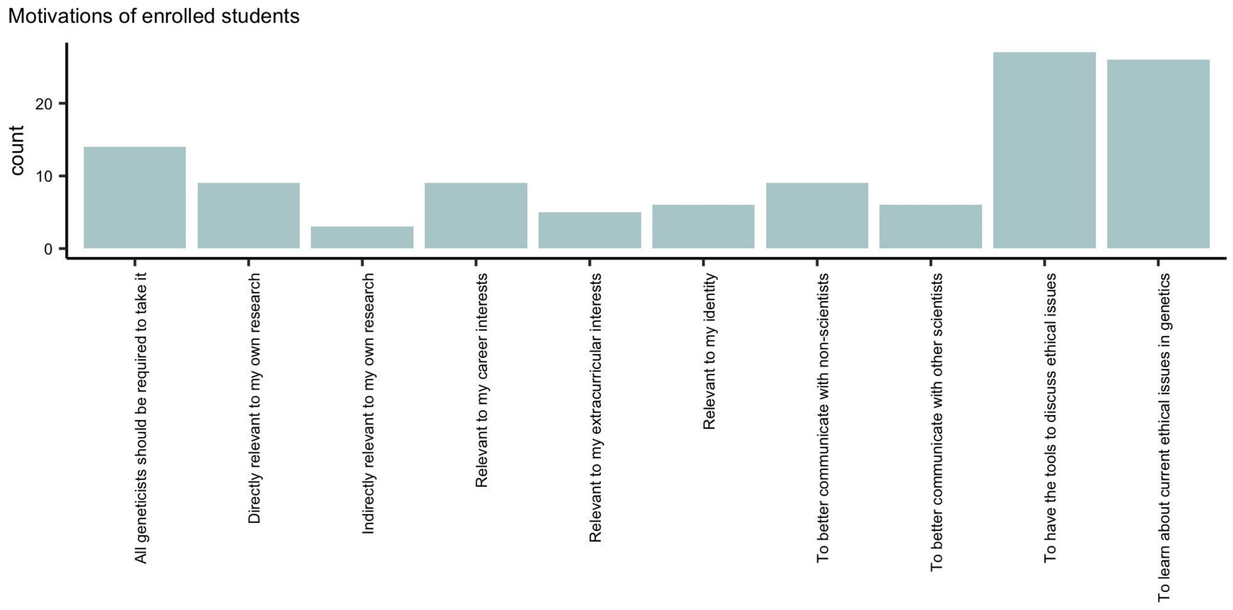


**Supplemental Figure 2: Motivations of enrolled students.** Students were asked to rank the relative importance of 10 potential motivations for taking the course. The histogram depicts the number of students that cited a given motivation as one of their top 3 reasons for taking the course. Data points are aggregated between 2022 and 2023.

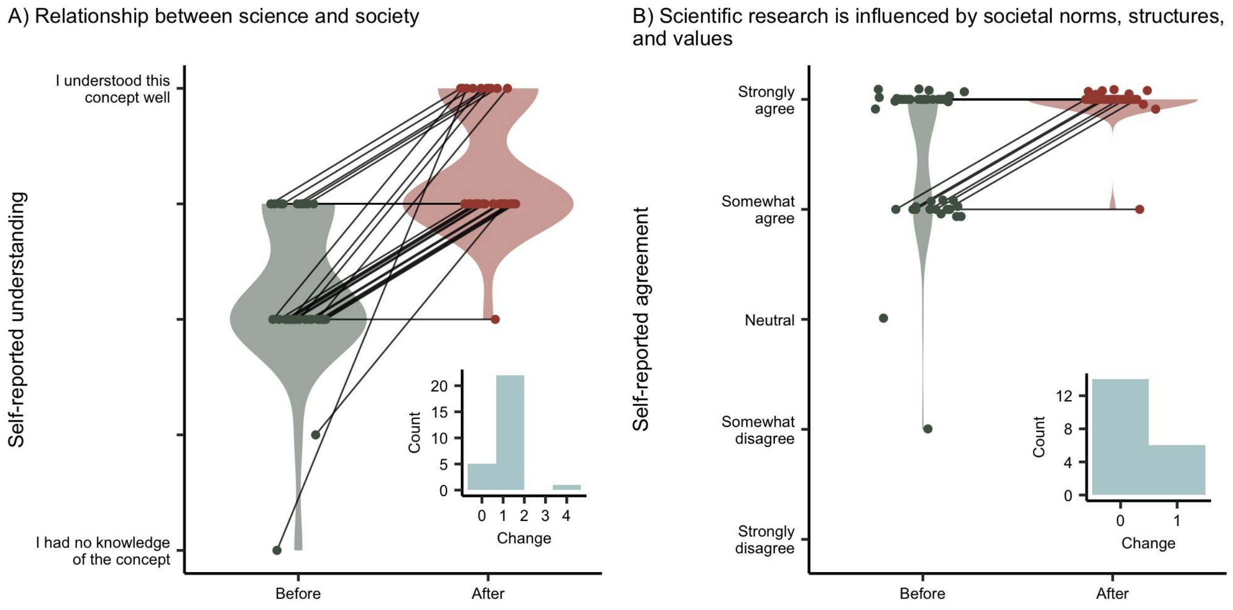


**Supplemental Figure 3**: **Perceptions of science and society in student evaluations.** A) Change in self-reported understanding of the "relationship between science and society" before and after the course. B) Change in self-reported agreement with "scientific research is influenced by societal norms, structures, and values" before and after the course. Data points are aggregated between 2022 and 2023 and jittered for enhanced visibility. Violin plots are included for illustrative purposes, with solid lines connecting the pre-course and post-course responses from the same student when applicable. Plot insets illustrate the change in opinion between pre-course and post-course responses for students who took both surveys, where a value of one represents a 1-unit difference in understanding or agreement.

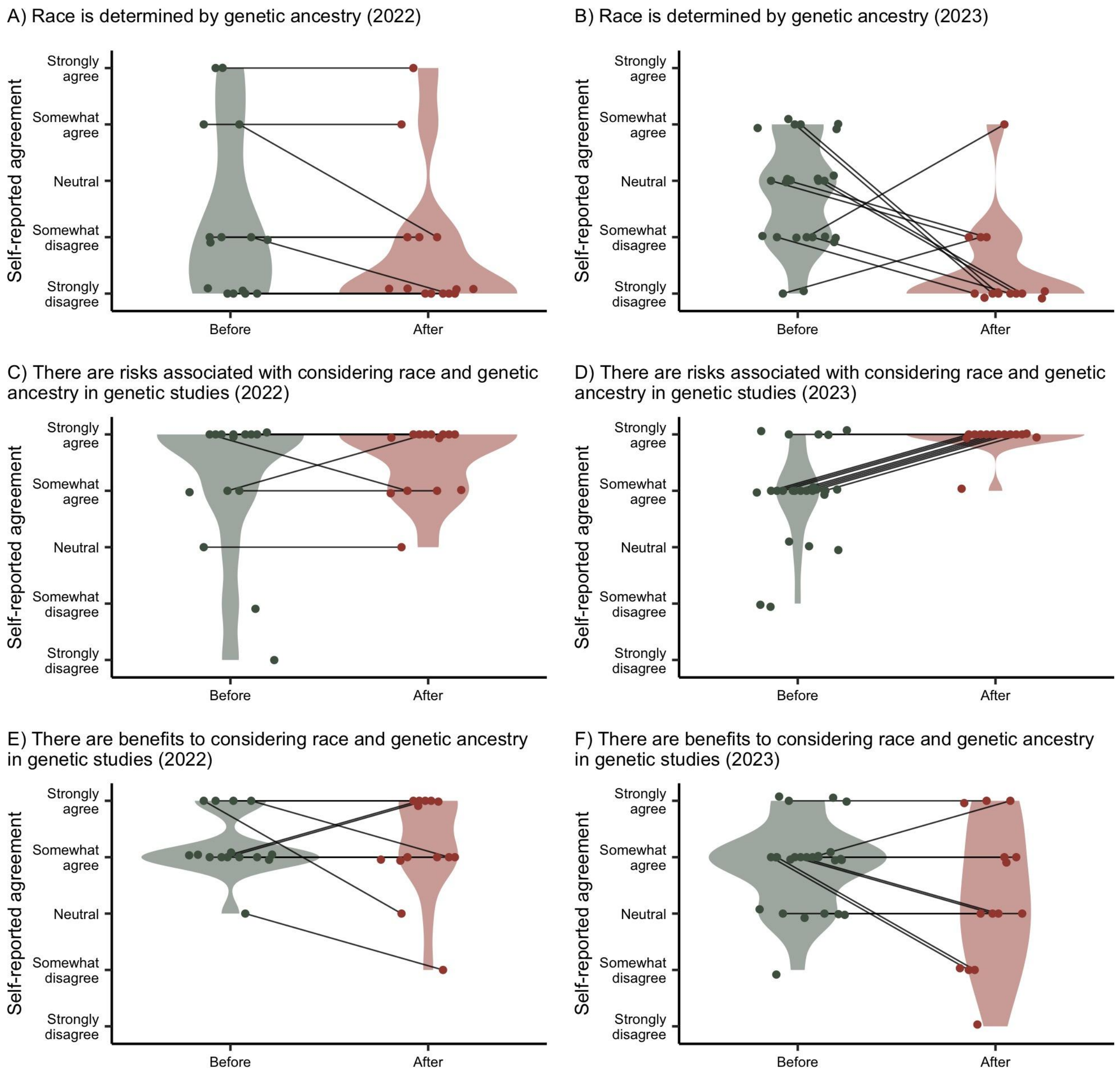


**Supplemental Figure 4: Perceptions of race, ancestry, and genetics in 2022 vs 2023 student evaluations.** A,B) Changes in self-reported agreement to “race is determined by genetic ancestry” before and after the course in 2022 and 2023. C,D) Changes in self-reported agreement to “there are risks associated with considering race and genetic ancestry in genetic studies” before and after the course in 2022 and 2023. E,F) Changes in self-reported agreement to “there are benefits to considering race and genetic ancestry in genetic studies” before and after the course in 2022 and 2023. Data points are jittered for enhanced visibility and violin plots are included for illustrative purposes, with solid lines connecting the pre-course and post-course responses from the same student when applicable.

### Supplemental Material: Stanford Archives Lesson Plan

#### Behavioral genetics, differences of sexual development, and criminality

##### Background

In January 1973 the Governor of California, Ronald Reagan, announced that the proposed Center for the Study and Reduction of Violence at UCLA was close to fruition. This program was staunchly opposed by many civil rights organizations, including the NAACP, the National Organization for Women (NOW), the Mexican-American Political Association, and the California Prisoners' Union. Part of the opposition to the Center was that it would focus on biological causes for aggressive behavior, rather than taking into account the social context/environment, and would attempt to falsely prove damaging stereotypes of marginalized individuals. For example, one proposed research focus of the center was understanding how XYY males might be more aggressive through mass screening programs. There were also fears around psychosurgery, or using surgical procedures to "correct" aggressive behavior, which had already gained a reputation for ethical abuses and exploitation of already-vulnerable populations. Thanks to a slew of public activism against the Center, the proposed state funding was eventually blocked, and the Center was never opened.

##### Themes

Genetic determinism, scientific activism, sex and gender, stakeholdership and community engagement, race and genetics

##### Documents

- Newspaper cutting: "Genes May Tell One's Potential to Commit Crimes"
- Newspaper article: "Proposed violence center 'swirling in controversy'"
- Newspaper article: "Merits of Center for the Study of Violence Debated"
- Daily Bruin article: "Vote YES against the Violence Center"
- LA Times article: "A Strife-Torn Violence Center"
- Letter: Special interest groups representing marginalized communities writing to advocate against this center
- Flier: Asks people to write to their state representative
- Letter: Thank you letter to California State Representative Willie Brown for preventing state funding of Violence Center

##### Discussion questions

- What were the reasons the UCLA Center for Study and Reduction against Violence was initially created?
- Who were the stakeholders in this research? Who opposed the Center, and why?
- What actions did activists take to work against the Center and what was the result?
- What do you think the broader impacts on society would have been if the Center had been funded?

- What connections do you see between this case example and other topics we've discussed in class?

##### Forced sterilization

###### Background

The eugenics movement of the 20th century had a global reach, with forced sterilization programs implemented in countries around the world. The movement reflected a belief in the genetic superiority of certain groups and the desire to eliminate those deemed "unfit" or "undesirable." These policies were deeply connected to racism, classism, ableism, and xenophobia, and resulted in the forced sterilization of thousands, if not millions, of people. The legacy of forced sterilization continues to impact marginalized communities, particularly women of color, who were disproportionately affected by these practices. Chicano-focused journalism informed their community about these sterilization efforts, and highlighted efforts of activists fighting against it. The documents in this section are written in both English and Spanish, and they discuss forced sterilization across multiple locations, including Los Angeles, California; Puerto Rico; and India.

###### Themes

Reproductive genetics, eugenics, scientific activism

###### Documents

- Newspaper article: "Esterilizaciones forzadas" (in Spanish)
- Newspaper article: "The struggle against forced sterilization"
- Newspaper article: "Forced sterilization of third world women"
- Newsletter: "Sterilization: US Alternative to Liberation in the 3rd World"

###### Discussion questions

- What efforts did activists take against sterilization efforts?
- How do the global sterilization efforts discussed in these documents connect to race, class, and colonialism/imperialism?
- What is your reaction to seeing the discussion of sterilizations in these community-driven newspapers? What can we learn from this today?

##### Race, genetics, and IQ

###### Background

In 1916, Lewis Terman, a professor of psychology at Stanford, developed the Stanford-Binet intelligence test, forming the basis for modern-day IQ testing. Terman's work played a huge role in the eugenics movement in the early 20th century. Following the Holocaust, eugenics concepts fell out of favor for some time, but saw a resurgence in the mid 20th century thanks to William Shockley. Shockley was a Nobel laureate in physics and Stanford professor who spoke against improvements in education for African Americans, arguing that they were genetically less capable of education. Shockley later funded the work of Arthur Jensen, who would go on to be an educational psychologist at UC Berkeley and whose research claimed that genetic variation drove racial differences in IQ. Jensen was a major player in the rise of genetic determinism and scientific racism towards the end of the 20th century.

The materials in this section reference a variety of perspectives from key academics in this history – both those supporting racial essentialism and genetic determinism of IQ (Lewis Terman, William Shockley, Arthur Jensen) and those opposed (Stephen J Gould, Richard Lewontin, Leon Kamin, Luca Cavalli-Sforza, Herbert Aptheker).

###### Themes

Racial essentialism, genetic determinism, scientific activism

###### Documents

- Paper by Leon Kamin: "Heredity, Intelligence, Politics, and Psychology"
- Article: "Intelligence of Negro recruits"
- Stanford Daily article: Shockley v Cavalli Debate
- Stephen Jay Gould essay: discusses objectivity of science and scientific racism
- Personal correspondence: from John Cirace to Stephen Jay Gould demonstrating contradictory views on genetic determinism

###### Discussion questions

- Can you summarize the positions held by the pro-racial essentialism, pro-genetic determinism crowd (in these documents: Terman, Shockley, Jensen) and their opponents?
- What mechanisms did scientists use to push back against racist interpretations of science?
- Based on these documents, which social structures, norms, or values have influenced academics' interpretation of scientific results, and how have they done so?
- What connections do you see between the disagreements outlined in these documents and what we discussed in class session 5 about modern-day debates regarding genetic research on educational attainment?
- What is your perspective on modern-day genetic research on educational attainment? Has it changed after viewing these documents – and if so, how?

##### Scientific activism and social responsibility

###### Background

In the 1970s and 80s, in response to a divisive political climate and the resurgence of racism and sexism – both within the academic realm of evolutionary biology, as well as the Boston community and U.S. more broadly – two important left-wing organizations were formed at Harvard. The Committee Against Racism (CAR) was an organization dedicated to anti-racism and was one chapter of what would eventually become an international organization. CAR was active in protesting racist rallies by the KKK and other white supremacist groups, as well as promoting racial integration of the Boston public school system via "busing". In parallel, the

Sociobiology Study Group was an academic organization that focused on countering what they saw as the inherent racism and sexism in sociobiology, a sub-field that sought to explain various features of human society using the principles of evolutionary biology.

Prominent members of CAR and the Sociobiology Study Group included Stephen Jay Gould and Richard Lewontin (who wrote the iconic 1972 paper on “The Apportionment of Human Diversity”, declaring race to be of “virtually no genetic or taxonomic significance”).

###### Themes

Scientific activism, social responsibility, racial essentialism, genetic determinism

###### Documents

- Harvard Crimson article: “Laying the Foundation for a Racist Synthesis”
- Committee Against Racism Statement and Pamphlets: promoting busing for integration
- Meeting Notes: Sociobiology Study Group
- Sociobiology Study Group Workshop Agenda: “A Critique of Biological Determinism”
- Charles Coulston Gillespie essay: essay on the relationship between the scientist and the state with annotations from Stephen Jay Gould

###### Discussion questions

- What were the Sociobiology Study Group and Committee Against Racism working to fight? What actions did they take?
- How does this connect to contemporary issues? How might they be different or similar to modern efforts?
- Do you think these groups were successful? Why or why not?
- Stephen J Gould, a member of the Sociobiology Study Group and Committee Against Racism, wrote: “What have we a right to expect of a scientist?” What do you think we should expect of scientists in regards to:
    - personal biases and prejudices
    - relationship with the government/state
    - relationship with society

##### Eugenics and anti-miscegenation

###### Background

The American eugenics movement emerged in the late 19th and early 20th centuries, largely in response to Darwin’s theory of evolution and driven by the idea of improving the human species through selective breeding and the suppression of traits deemed undesirable. One of the key focuses of the movement was opposition to interracial marriage, as eugenicists believed that such unions would result in the "degeneration" of the white race. Many eugenicists were influential geneticists who argued that certain races were genetically inferior to others – in particular that Black people who were believed to be less intelligent than white people. They used their scientific authority to promote various laws and policies, including “anti-miscegenation” laws that criminalized interracial marriage. Materials in this section include views from both eugenicists, as well as those who opposed eugenics.

##### Themes

Eugenics, race and ancestry, racial essentialism

##### Documents

- Hand-drawn Family Tree: David Starr Jordan, "Chart of Fitness" capturing Jordan’s purported genealogy
- 1950s Era Pamphlet: “The Races of Mankind”
- Booklet: “The New Family and Race Improvement”
- Agendas: Second and Third International Eugenics Congress
- Court Case Transcript: “Miscegenation Law, Court Cases, and Ideologies of ‘Race’”
- Article: “I do not believe there is a superior race”
- Article: “Intermarriage and the race problem”

##### Discussion questions

- Who is included on David Starr Jordan’s family tree? Who might be excluded? Why do you think he created this document?
- What beliefs or assumptions about race, ancestry, and genetics can you identify in the arguments presented in the miscegenation law court case? How are they similar or different to how these concepts are perceived by society in the modern day?
- What topics were discussed at the eugenics congresses? What influence or impact do you think these “pseudo-academic” conferences had on society more broadly?
- What social or societal factors influenced individuals who supported eugenics and/or opposed interracial marriage? How were scientific ideas/concepts used to defend or justify their positions?
